# *Leptospira* seroprevalence in dogs, cats and horses: the effect of cross-reactivity between serovars and vaccination status on seroprevalence estimates

**DOI:** 10.1101/2024.04.17.589957

**Authors:** Kellie A. McCreight, Liana N. Barbosa, Odoi Agricola, Porsha Reed, Sreekumari Rajeev

**Affiliations:** Department of Biomedical and Diagnostic Sciences, College of Veterinary medicine, University of Tennessee, Knoxville

## Abstract

In this study, we estimated the *Leptospira* seroprevalence in dogs (n = 376), horses (n = 88), and cats (n = 169) using Microscopic Agglutination Test (MAT) against 12 *Leptospira* serovars. We observed a *Leptospira* seroprevalence of 29.41%, 47.73%, and 12.35% in dogs, horses, and cats, respectively. The highest seroprevalence was observed for serovar Autumnalis (74.55%) in dogs, and Bratislava in horses (95.24%) and cats (42.86%). We found a significant level of potential cross-reactivity between multiple *Leptospira* serovars tested, with highest cross-reactivity to serovar Autumnalis in dogs. The dogs that were vaccinated against *Leptospira* had a significantly higher seroprevalence (45.92%) compared to unvaccinated dogs (16.28%; p < 0.001). A significant difference in seroprevalence was observed in vaccinated and unvaccinated dogs to all four serovars included in the canine vaccine (p < 0.001). Among the diagnostic samples submitted from 2021-2023, 40% (103/252) of the canine serum samples were positive by MAT, with the highest positivity rate for the serovar Autumnalis. On *Leptospira* PCR, 10.7 % (35/325) urine samples and 5.8% (15/257) blood samples were positive. Our study findings show evidence of *Leptospira* exposure and potential disease in the study area. The cross-reactivity between the *Leptospira* serovars used in the MAT and vaccination status may also overestimate the levels of exposure.

## Introduction

Leptospirosis is a spirochete bacterial infection and a cause of life-threatening disease in humans and animals [1, 2]. It is a re-emerging zoonosis resulting in approximately 1 million new human cases and over 50,000 deaths worldwide [3]. *Leptospira* infection in domestic animals such as dogs, cats, horses, and cattle may lead to fatal disease or reservoir status, however, accurate estimates of animal infections and its impact are not available [2]. Many mammalian hosts, including the abovementioned domestic animal species, may act as reservoir hosts by harboring *Leptospira* in their kidneys and reproductive tract and can continuously excrete the bacteria through urine and thereby contaminating the environment. New infection occurs from direct contact with infected animal’s urine and from contact with the contaminated environment. Animal leptospirosis is well-recognized and clinical symptoms may range from mild febrile illness to multiorgan failure and death, or other reproductive diseases in livestock species. However, the actual disease incidence is still not documented due to the lack of awareness and challenges involved in getting an accurate diagnosis. A definitive disease diagnosis can only be achieved by the detection of the infecting *Leptospira* strain by PCR or culture in a patient with compatible clinical signs. The Microscopic agglutination test (MAT) is a widely used method for detecting serovar specific *Leptospira* antibodies, and to identify exposure to circulating serovars in a particular geographic region. In humans, a single MAT titer of 800- or four-fold difference in acute vs. convalescent titer, is considered for definitive diagnosis. However, interpretation of MAT titer can be confusing in case of animals due to potential widespread exposure, vaccination status, and asymptomatic infection. The infection prevalence and the infecting/circulating serovar vs. species may vary geographically and seasonally and hence frequent updating of prevalence data is highly desirable.

A recent regional study conducted in the Cumberland Gap region (CGR) close to the intersection of Kentucky, Tennessee, and Virginia [4], found that 13.13% of the dogs tested were positive for *Leptospira* DNA in urine by qPCR and 18% of the animals were seropositive. Nineteen of these dogs were from shelters in Tennessee of which only one dog was positive for MAT, and none were positive by PCR. There have been no equine studies reported in Tennessee. Based on only one previous study available on the seroprevalence of *Leptospira* in cats in Tennessee, all 50 cats in that study tested negative by both qPCR and MAT [5]. We have diagnosed many canine clinical leptospirosis cases in our diagnostic laboratory. Emerging zoonotic diseases such as leptospirosis are a public health concern, due to the potential of animals that can become reservoirs and carriers and contaminating the surrounding environment. A recent study we conducted using metagenomic sequencing and analysis identified a diverse group of pathogenic *Leptospira* present in the water and soil in the vicinity [6]. Therefore, to improve our understanding of extent of *Leptospira* exposure, we investigated the *Leptospira* seroprevalence in dogs, cats, and horses in Tennessee.

## Materials and methods

### Samples

We used clinical samples submitted for general testing from dogs, cats, and horses collected between January and August 2022 from the University of Tennessee, College of Veterinary Medicine, Clinical Pathology Diagnostic Laboratory. These samples are considered as biowaste and discarded after required testing. For this study, we obtained 374 canine samples, 169 feline samples, and 88 equine samples and stored these samples at -20°C until use. In addition, we also evaluated the results from canine *Leptospira* diagnostic submissions for the years 2021-2023.

### Microscopic Agglutination Test (MAT)

All serum samples were tested for the presence of antibodies against twelve *Leptospira* serovars, using the standard operating procedure from our *Leptospira* diagnostic and research laboratory. Briefly, 4-7-day old cultures of *Leptospira* serovars, maintained in the laboratory through continuous passage in Ellinghausen-McCullough-Johnson-Harris (EMJH) medium at 28°C, were used for MAT. The bacterial concentration was adjusted at a transmittance value between 75% and 80% on the spectrophotometer. Serum samples (50µl) at 1:25 dilution in Phosphate Buffered Saline (PBS) was treated with 50µL of each serovar in a flat bottom 96 well microtiter plate and then incubated for 1.5-2 hours at 28°C. Homologous antisera prepared in rabbits for each serovar used in the assay served as the positive control and PBS as the negative control. The plates were read using a dark-field microscope, using a 5x long working distance objective. The samples with 50% agglutination were considered positive at 1:50 final dilution. Antibody titer was determined for all the samples showing more than 50% agglutination.

### Data analysis

IBM SPSS Statistics (Version 27 and 28, IBM, New York) was used to estimate prevalence. A 2-tailed Spearman rank-order correlation test was performed for the 12 serovars used in the study to determine the potential cross-reactivity between serovars. In dogs, a comparison of seroprevalence between the vaccinated positive and unvaccinated positive groups as well as the association between serovars included in the vaccine were done using Fishers exact test. A p-value below 0.05 is considered statistically significant.

## Results

### *Leptospira* seroprevalence in dogs, cats and horses

Overall seroprevalence of 29.41%, 47.73% and 12.35% was observed in dogs, horses and cats, respectively (Fig 1). We observed detectable agglutinating antibody response to 10 out of 12 serovars tested in dogs. No antibody response to serovars Bataviae and Tarassovi was observed. Among the serovars tested in dogs, the highest seroprevalence was observed for the serovar Autumnalis (82/110; 74.55%) followed by Grippotyphosa (44/110; 40.0%), and the lowest seroprevalence was observed for serovar Hardjo (3/110; 2.73%). The MAT titers ranged from 1:50 to 1:1600 in canine samples. The highest titers were observed in Bratislava (1:1600) followed by Autumnalis (1:800), Grippotyphosa (1:800) and Pomona (1:800). Titer distribution to each serovar in dogs is shown in Fig 2. In horses, all the serovars tested had detectable agglutinating antibody response. Among the serovars tested, the highest seroprevalence was observed for Bratislava (40/42; 95.24%), followed by Copenhageni (24/42; 57.14%), and the lowest seroprevalence was observed for Tarassovi (1/42; 2.38%). The highest titer in horses was observed for both Bratislava and Pomona (1:400). Titer distribution to each serovars in horses is shown in Fig 3. In cats, the highest seroprevalence was observed in Bratislava (9/21; 42.86%) followed by Hardjo (8/21; 38.10%), and the lowest seroprevalence was observed for serovars Bataviae (1/21; 4.76%) and Mankarso (1/21; 4.76%). Seroprevalence to Ballum and Tarassovi were not observed. The antibody titers ranged from 1:50 to 1:3200. The highest titer was observed for the serovar Hardjo (1:3200), however, the majority of the feline samples only had a 1:50 titer. Titer distribution to each serovars in cats is shown in Fig 4.

**Figure 1.**
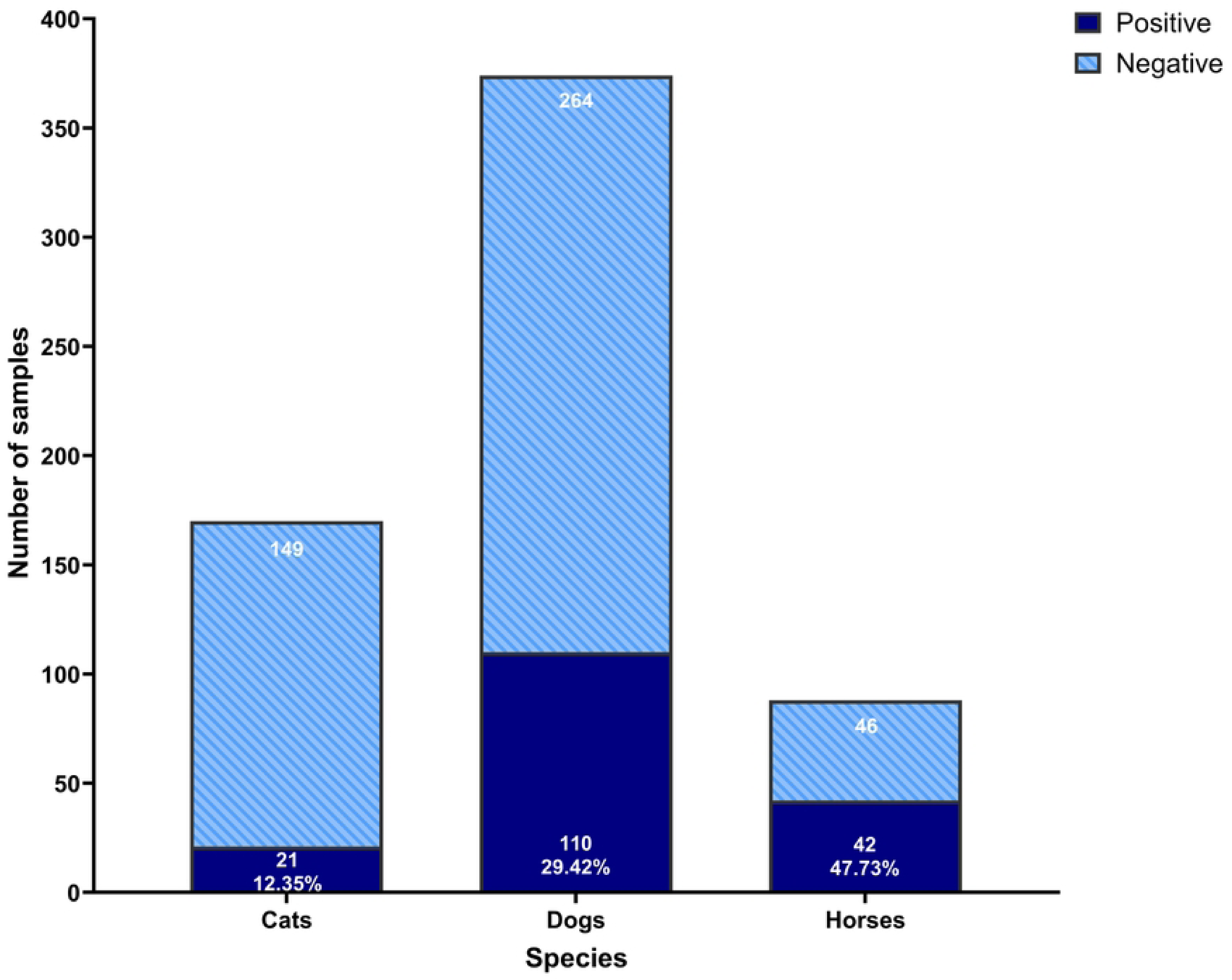
Overall *Leptospira* seroprevalence detected by screening serum with MAT in cats, dogs and horses.

**Figure 2.**
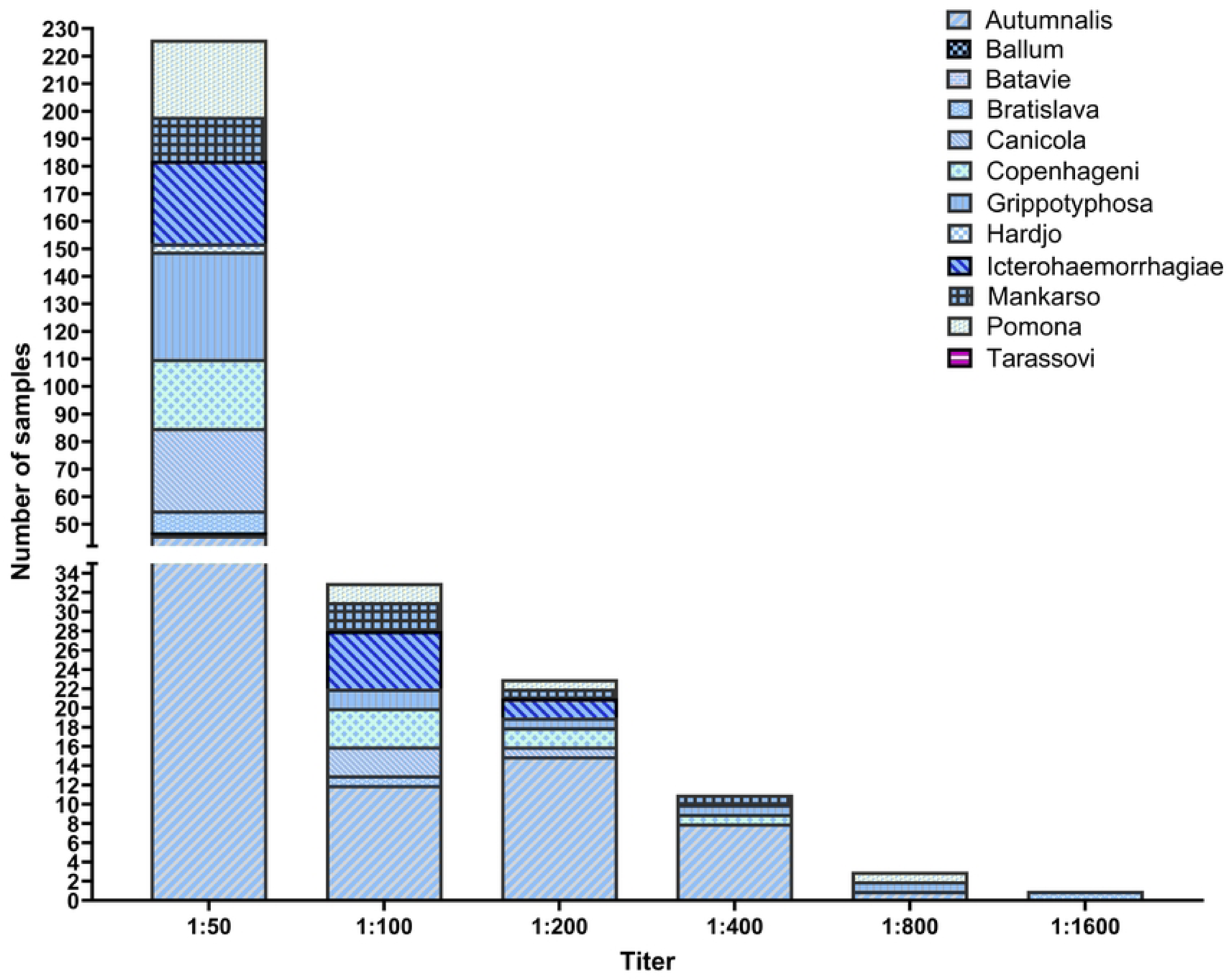
MAT Titer ranges to *Leptospira* serovars tested in dogs.

**Figure 3.**
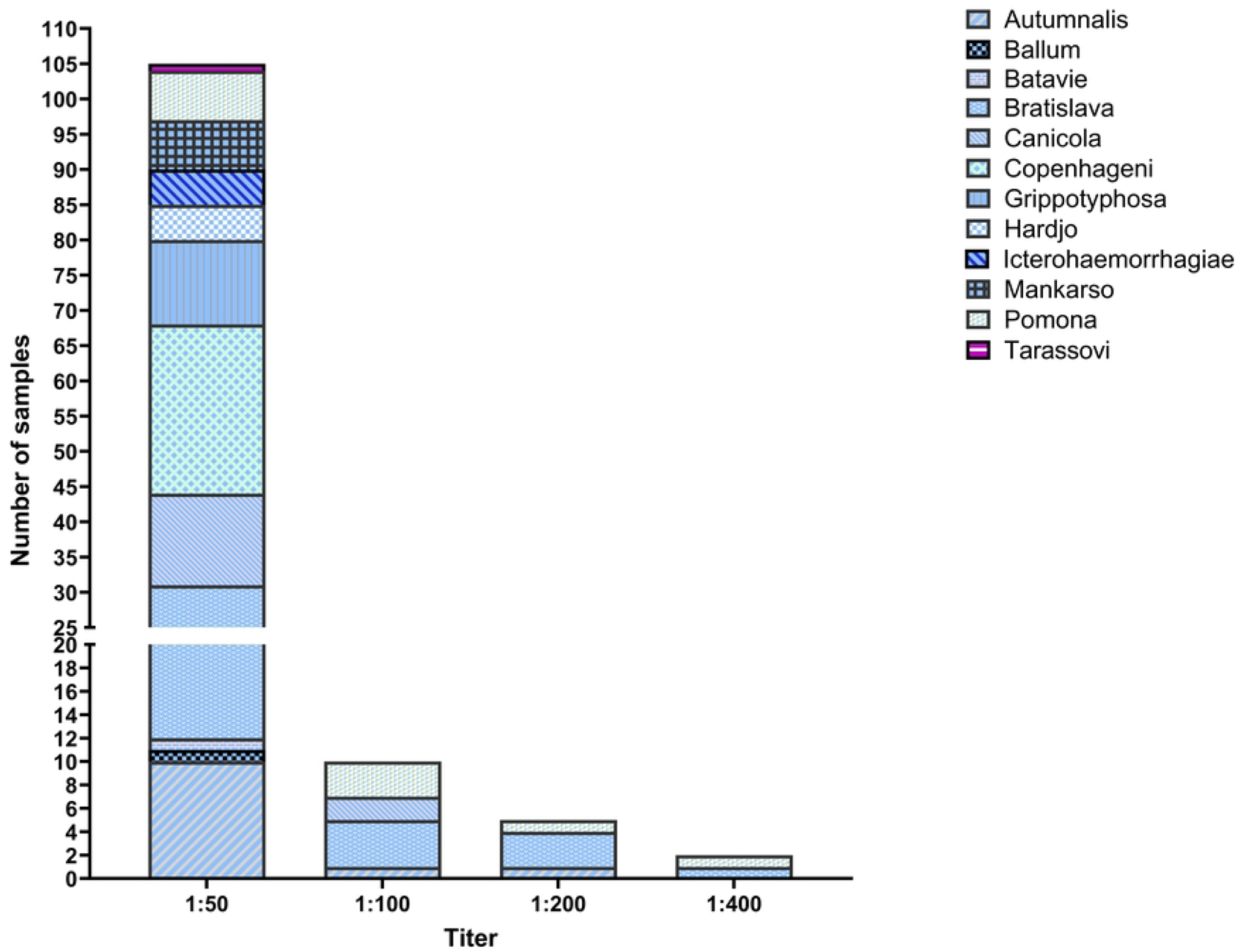
MAT Titer ranges to *Leptospira* serovars tested in horses.

**Figure 4.**
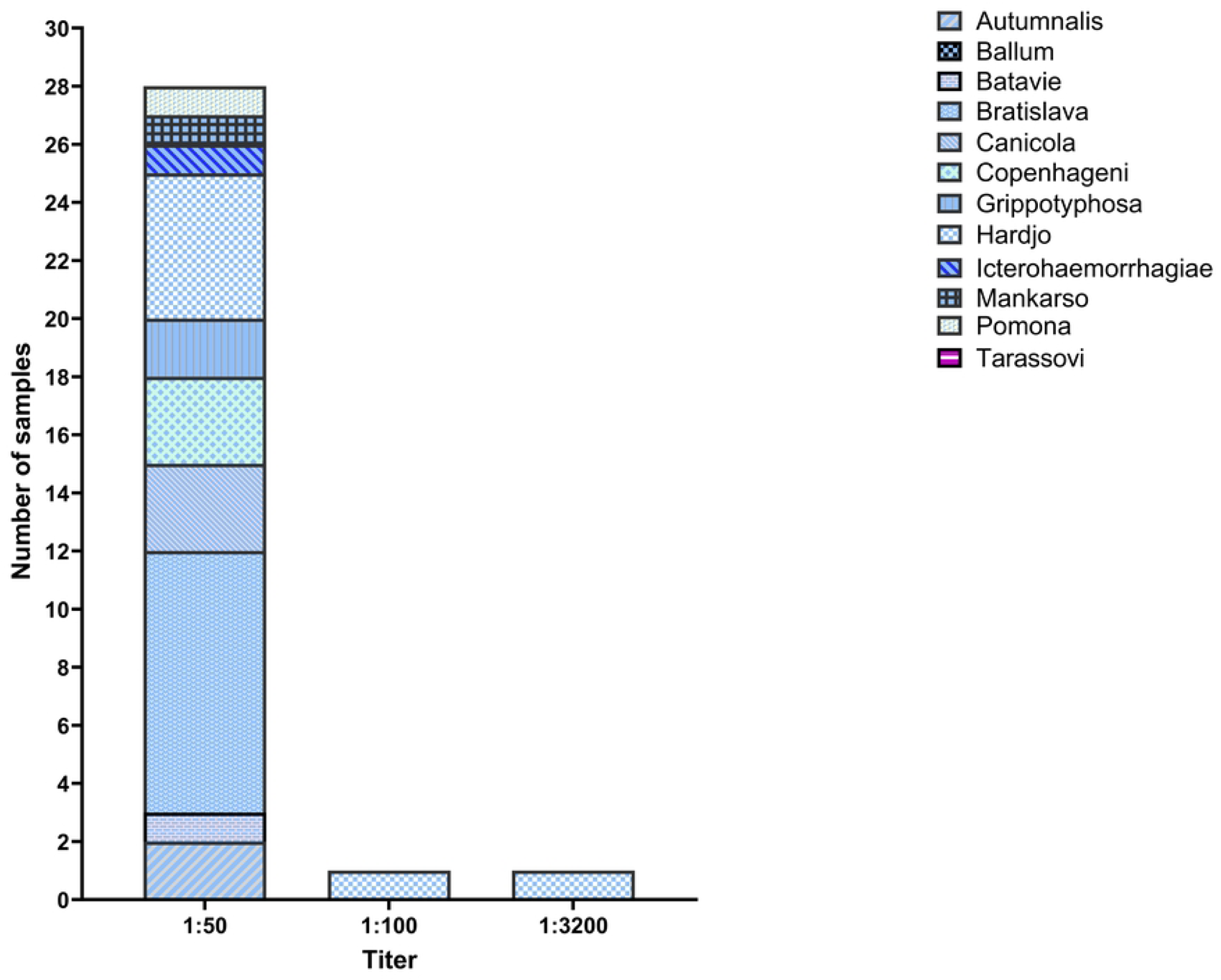
MAT Titer ranges to *Leptospira* serovars tested in cats.

### Age, sex, breed and County distribution

We also evaluated age, sex, and breed distribution, and the geographic information related to the samples tested. The majority of the samples came from dogs in the 6-year to 10-year age range. Of the 374 samples, 191 (51.07%) samples were from female dogs and 183 (48.93%) were from male dogs. For breed distribution, the population was overrepresented by mixed breed (97/374; 25.9%). Out of the 12 dog breeds identified, hunting dogs had the highest positivity rate (6/12; 50.0%), followed by working dogs and small breed dogs (3/12; 25.0% each). For cats, most samples were in the 6-year to 10-year age range. Of the 170 samples, 78 (45.88%) were from females, 91 (53.53%) were from males, and 1 (0.59%) was unknown. For breed distribution, the population was overrepresented by Domestic Shorthair (126/170; 74.18%). For horses, the majority of the samples were in the 0–5-year age range, with the least number of samples in the > 30-year range. Of the 88 samples, 26 (29.55%) were from females, 61 (69.32%) were from males and 1 (0.01%) was unknown. The breed distribution was overrepresented by American Quarter Horse (20/88; 22.73%). In this study we were able to obtain at least one sample from 25/95 (26.32%) of the counties in Tennessee. There are 4 regions in Tennessee, with most of the samples collected from region 1. For the canine samples, we had 304 (81.28%) samples from region 1, 59 (15.78%) from region 2 and 11 (2.94%) from region 3. We had no samples from region 4. For cats, we had 145 samples (85.29%) from region 1, 14 (8.24%) from region 2, 11 (6.47%) from region 3 and no samples from region 4. For horses, we had 65 samples (73.86%) from region 1, 18 (20.45%) from region 2, 4 (4.55%) from region 3, no samples from region 4. One sample (1.1%) did not have a county listed. The age, sex, breed, and geographic distribution data are summarized in S1 Appendix.

### Occurrence of potential cross-reactivity between serovars in MAT

Since many of the canine serum samples had positive reactivity to multiple serovars tested, we examined whether this could be due to cross-reactivity between *Leptospira* serovars. A Spearman rank-order correlation assessment suggested potential cross-reactivity between samples (Table 1). The serovar Autumnalis had the highest cross-reactivity among all other serovars reacting with 7 other serovars. A Venn diagram showing the reactivity between serovars is shown in S1 Fig. A similar observation was made for horses (data not shown). There were not enough positive samples for a Spearman rank-order correlation to be performed for cats.

**Table 1.** Spearman rank correlations between MAT positivity to different *Leptospira* serovars in dogs.

| Serovars | Autumnalis | Bratislava | Canicola | Copenhageni | Grippotyphosa | Icterohaemorrhagiae | Mankarso | Pomona |
| --- | --- | --- | --- | --- | --- | --- | --- | --- |
| Autumnalis | 1.00 |  |  |  |  |  |  |  |
| Bratislava | 0.265 | 1.00 |  |  |  |  |  |  |
| Canicola# | 0.505 | 0.158 | 1.00 |  |  |  |  |  |
| Copenhageni* | 0.625 | 0.359 | 0.591 | 1.00 |  |  |  |  |
| Grippotyphosa# | 0.584 | 0.190 | 0.416 | 0.552 | 1.00 |  |  |  |
| Icterohaemorrhagiae*# | 0.397 | 0.238 | 0.398 | 0.645 | 0.414 | 1.00 |  |  |
| Mankarso* | 0.540 | 0.323 | 0.520 | 0.818 | 0.529 | 0.642 | 1.00 |  |
| Pomona# | 0.731 | 0.203 | 0.447 | 0.608 | 0.539 | 0.452 | 0.596 | 1.00 |

All the correlations were statistically significant ( $p < 0.001$ ). (\*) Serovars belong to the same serogroup. (#) Serovars present in the vaccine. Red fonts are the serovars with moderate to strong correlation coefficient ( $> 0.4$ ).

### Comparison of seroprevalence between vaccinated and unvaccinated dogs

Since *Leptospira* vaccination is practiced in dogs, we evaluated the difference in seroprevalence in vaccinated vs. unvaccinated canine population. We separated the MAT results from samples for vaccinated and unvaccinated patients using clinical history obtained from hospital records. The vaccination data was available for 184 of the 374 canine patients. Of the 184 serum samples collected, 98 (53.26%) had been from vaccinated dogs and 86 (46.74%) from unvaccinated dogs. Of the 98 samples from vaccinated dogs, 53 (54.08%) were MAT negative for antibodies and 45 (45.92%) were MAT positive to one or more *Leptospira* serovars tested. The 86 samples from unvaccinated dogs were also further categorized as MAT positive or negative samples, with 72 (83.72%) testing negative and 14 (16.28%) testing positive. The vaccinated dogs had a significantly higher seroprevalence compared to the unvaccinated dogs (Table 2). The dogs that were vaccinated and tested positive, the most common serovars with MAT reactivity were Autumnalis (36/45; 80.0%), followed by Grippotyphosa (22/45; 48.89%), Icterohaemorrhagiae (20/45; 44.44%) and Canicola (18/45; 40.0%). When we tested the difference in seroprevalence to the 4 serovars present in the canine vaccine, a significant difference was observed for all the serovars included in the vaccine; Canicola, Grippotyphosa, Icterohaemorrhagiae and Pomona (p < 0.001) between the vaccinated and unvaccinated dogs (Table 3).

**Table 2.** Comparison of seropositivity between vaccinated and unvaccinated dogs.

|  | Number of positive samples/Number of samples<br>(Seroprevalence) | 95% Confidence interval |
| --- | --- | --- |
| Vaccinated | 45/98<br>(45.92%) | 45.7 – 60.3 |
| Unvaccinated | 14/86<br>(16.28%) | 39.7 – 54.3 |

(%) Percentage.

**Table 3.** Comparison of seropositivity between vaccinated and unvaccinated dogs to vaccine serovars.

|  | Prevalence in vaccinated group (%)<br>[95% Confidence interval] | Prevalence in unvaccinated group (%)<br>[95% Confidence interval] |
| --- | --- | --- |
| Canicola | 18.37<br>[0.038-0.289] | 3.49<br>[0-0.255] |
| Grippotyphosa | 22.68<br>[-0.098-.0131] | 5.81<br>[0-0.103] |
| Icterohaemorrhagiae | 20.41<br>[-0.082-0.191] | 4.65<br>[0-0.174] |
| Pomona | 14.29<br>[-0.005-0.213] | 6.98<br>[0-0.190] |

(%) Percentage.

### *Leptospira* testing results from the diagnostic tests

Our *Leptospira* Diagnostic and Research Laboratory offers PCR and MAT, or a panel of both to our veterinary clients. We performed diagnostic tests on 833 samples from canine patients submitted for *Leptospira* testing (MAT and/or PCR) from January 2021-December 2023. Forty percent (103/252) of the serum samples were positive by MAT with the highest positivity rate for the serovar Autumnalis (Fig 5). Approximately 10.7 % (35/325) of urine samples and 5.8% (15/257) of blood samples were positive by *Leptospira* PCR.

**Figure 5.**
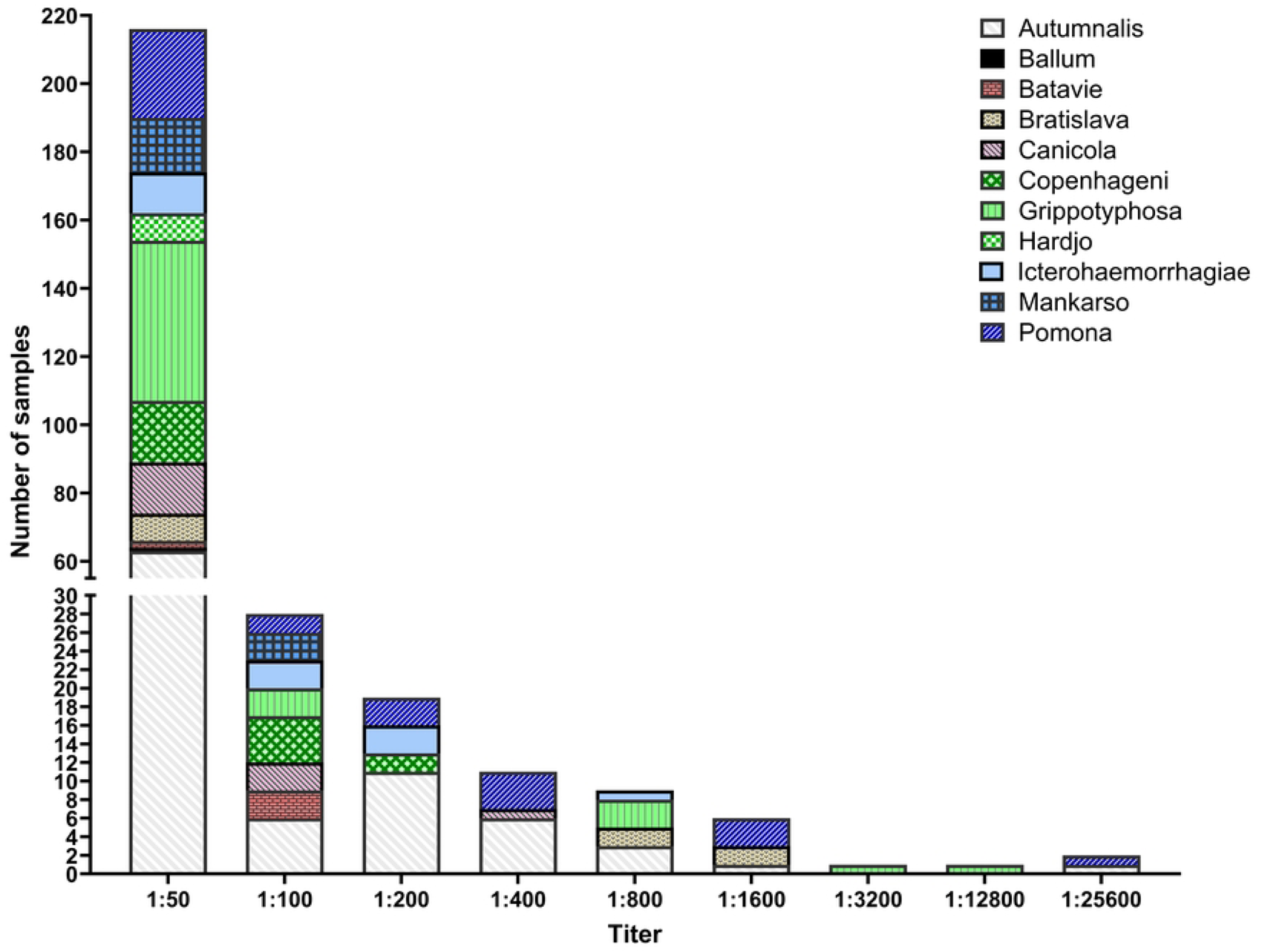
MAT titer range to *Leptospira* serovars in diagnostic samples from dogs.

## Discussion

The overall prevalence of *Leptospira* exposure is widely under-documented in animals and many of the factors influencing the seroprevalence rates are not known. Our seroprevalence study updates the antibody prevalence to *Leptospira* in dogs, cats and horses in Tennessee. We also report the factors that might influence the assessment of seroprevalence. Data on *Leptospira* infection in dogs range from those studies evaluating the seasonal prevalence patterns, to spatial and spatio-temporal clustering, to seroprevalence throughout the United States [7–10]. The prevalence in dogs in the United States ranged from 4.48% to 12.03% from previous reports [9]. Only one previous study is available on the *Leptospira* seroprevalence in dogs and cats was conducted in a limited geographic region in Tennessee [5].

In our study the highest MAT reactivity in dogs was observed against serovar Autumnalis, followed by the serovars, Bratislava, Grippotyphosa and Pomona, and our findings were consistent with previous findings [11, 12]. The MAT, a widely used test in *Leptospira* seroprevalence studies, detects agglutinating antibodies to the lipopolysaccharide components of *Leptospira* serovars using a limited array of serovars potentially circulating in a geographic area of interest. The test uses live *Leptospira* cultures maintained continuously in the laboratory and has acceptable specificity since antibodies to *Leptospira* do not cross-react with other bacterial species. However, sensitivity of this test can be low when compared to other tests, especially in the initial stages of infection. The test is laborious, difficult to standardize and has varying levels of subjectivity in reading and interpretation of the results. The MAT positive results may typically indicate the presence of antibodies from current infection, previous exposure or recent vaccination against *Leptospira.* In routine MAT testing, a typical cutoff serum dilution is 1:100, but cutoff as low as 1:10 can be used for increasing the test sensitivity. In our lab we typically use a cutoff of 1:50. In this study, in addition to establishing *Leptospira* seroprevalence data in dogs, cats and horses in Tennessee using MAT, we identified multiple issues that may affect the outcome of MAT in surveillance and diagnostic testing. The utility of MAT is also limited based on the number of serovars that can be included in the panel. This test is often described as reference standard test diagnosing *Leptospira* infection in acute and convalescent cases[13]. In our experience in canine patients with initial stages of infection, there is little antibody response detected by MAT which can often confuse the clinician. Diagnosing clinical leptospirosis is the initial stages of infection in patients with clinical suspicion is critical for implementing intervention strategies to avoid further complications from disease. It should be emphasized that MAT results alone should not be used in these cases as interpretation can be difficult due to the low titer level observed in early infection. Based on the range of titers observed in this study a single titer at any level should be interpreted with caution. While this information can help practitioners deduce their conclusion combining other factors, they should be encouraged to conduct combination testing that includes PCR from blood and urine samples.

We identified several factors that could influence the interpretation of serologic data for *Leptospira* surveillance and diagnostics. One of the significant observations from our study is the shared seroreactivity observed to multiple serovars. This is often misinterpreted as true seroprevalence to multiple serovars in many studies. There was a significant correlation of seroreactivity between multiple serovars, specifically Autumnalis, Canicola, Copenhageni, Grippotyphosa, Mankarso and Pomona. Autumnalis is not a common serovar identified in dogs in the USA or elsewhere. It is noteworthy that some of the serovars that cross-reacted belong to the same serogroup, such as serovars Icterohemorrhagiae, Copenhageni and Mankarso belonging to serogroup “Icterohaemorrhagiae”, which may explain cross-reactivity between those serovars. Such results should be interpreted with caution to avoid overestimation of seroprevalence. In the initial stages of clinical leptospirosis, cross-reactivity to even unrelated serovars described as ‘paradoxical reactions’ is often observed. In our case, the shared MAT reactivity with Autumnalis with other serovars may be resulting from cross-reactivity to surface antigens common to these serovars.

Vaccination is commonly practiced in dogs using a quadrivalent bacterin vaccine that consists of serovars Canicola, Grippotyphosa, Icterohaemorrhagiae and Pomona. *Leptospira* vaccination status in dogs was only available for 49% of the samples collected, and *Leptospira* vaccination status in horses was not available to us, so we could not assess potential effect of vaccination in horses. The presence of serovars in vaccine formulations may also lead to MAT cross-reactivity in vaccinated animals, as shown in our data. The significant difference observed between vaccinated and unvaccinated dogs to serovars Canicola, Grippotyphosa, Icterohaemorrhagiae and Pomona are thought to be related to the vaccine response. The serovar Autumnalis is not included in the canine vaccine, but the positivity and cross-reactivity observed in this group to this serovar was significantly higher. A pattern consistent with the seroprevalence data from the random canine population was noticed in diagnostic samples showing highest seroprevalence to the serovar Autumnalis. The highest titers consistent with clinical leptospirosis were observed against serovars Grippotyphosa, Pomona and Autumnalis. Many of the diagnostic samples positive by PCR, when sequenced were related to *L. kirschneri* serovar Grippotyphosa. *Leptospira* infection is not directly linked to any specific breed of animal, but literature suggests that for dogs, small breeds and hunting breeds could become infected more often than others [14–17]. Out of the 12 dog breeds identified in our study, half of the positive samples came from hunting dogs, working dogs and small breed dogs.

Among the three animal species tested, the highest prevalence was observed in horses. Equine studies are available and range from serologic evidence or diagnosis of leptospirosis in the United States [18, 19] to challenges of establishing experimental *Leptospira* infection [20]. The prevalence in horses in the United States ranged from 2.5% to 16.4% is lower than our study [21, 22]. In horses, the highest seroprevalence was with serovars Bratislava, Copenhageni and Canicola. Serovars Bratislava and Canicola have shown seropositivity in horses in previous studies in the United States [23, 24], agreeing with our findings. No studies in the United States have been done to assess the risk of equine breeds and *Leptospira* infection. In horses, a bacterin vaccine containing *L. interrogans* serovar Pomona is available, but we could not retrieve reliable vaccination history for this species to include in our analysis.

Feline studies range from literature reviews aiding in diagnoses [25], to shedding, seropositivity, and seroprevalence [5, 26]. The prevalence in cats in the United States ranges from 4% to 33% [25]. Based on only one previous study available of the prevalence of *Leptospira* in cats in Tennessee, and all the cats in that study tested negative [5]. In our study, the serovars that had the highest number of positive samples were serovars Bratislava and Hardjo, and the highest titer seen in cats was against serovar Hardjo (1:3200). Positivity to serovar Bratislava agrees with other feline studies both inside and outside of the United States [26–29], while a limited number of studies reported positive samples against serovar Hardjo [25, 27].

One of the limitations of our study was that most samples were from East Tennessee and a small number of samples from the other regions making it difficult to assess the prevalence in these regions accurately. In addition, we used a set of convenient serum samples submitted for other evaluations, not specifically for *Leptospira* testing and therefore the random sampling might have decreased the possibility of selection bias, and potentially provided an equal chance of selection. However, it is disadvantageous for its inability to guarantee that the data collected is reflective of the entire population. However, the prevalence data from our diagnostic samples targeting only samples from animals suspected to have leptospirosis was similar to the results from the random samples.

In conclusion, we estimated *Leptospira* seroprevalence in dogs, horses and cats in our geographic region and we conclude that the cross-reactivity between the *Leptospira* serovars used in the MAT can potentially overestimate serovar level seroprevalence. In addition, vaccination data, if not included in the analysis, may also overestimate the actual prevalence of infection. Future research should focus on the significant associations found in this study and pair the MAT with a highly specific test, such as Real-Time PCR, to make an early and actionable diagnosis for effective intervention. In one of our recent studies a high diversity of *Leptospira* was identified in water and soil samples in the region emphasizing the potential for getting infection from these sources [6]. *Leptospira* culture and characterization to identify the species and serovars from animals with clinical leptospirosis should be attempted and it will improve our understanding of epidemiology and ecology of this serious zoonotic disease. In addition, better understanding of reservoirs, transmission mode and impact on other animals is also needed. Accurate estimates of *Leptospira* infections in animals and its health impact are not available. Future studies must focus on the variations clinical and pathologic effects and its impact following *Leptospira* infection in animals.

## Acknowledgements

We would like to thank UTCVM Clinical Pathology and Bacteriology and Mycology diagnostic laboratory staff for their support in collecting clinical samples.

## Supporting information

**S1 Appendix. Age, breed, sex and geographic distribution data.** (A) Age distribution of 374 canine samples tested. (B) Sex distribution of 374 canine samples tested. (C) A map of regions I-IV in Tennessee presented by the Tennessee County Highway Officials Association (TCHOA). (D) Total number of samples and total number of positive samples by Tennessee region in dogs. (E) Canine breed distribution. (F) Age distribution of 88 equine samples tested. (G) Sex distribution of 88 equine samples tested. (H) Total number of samples and total number of positive samples by Tennessee region in horses. (I) Age distribution of the 170 feline samples tested. (J) Sex distribution of the 170 feline samples tested. (K) Total number of samples and total number of positive samples by Tennessee region in cats. (.DOCX)

**S1 Fig. Venn diagram showing the 6 highest cross-reacting Leptospira serovars shown in relation to the 9 other serovars.** (A) Autumnalis. (B) Canicola. (C) Copenhageni. (D) Grippotyphosa. (E) Mankarso. (F) Pomona. (.DOCX)

